# A New Mouse Model of Diffuse Midline Glioma to Test Targeted Immunotherapies

**DOI:** 10.1101/2021.10.15.464284

**Authors:** M Seblani, M Zannikou, J.T. Duffy, R.N. Levine, Q Liu, C.M. Horbinski, O.J. Becher, I.V. Balyasnikova

**Affiliations:** Ann & Robert H. Lurie Children’s Hospital of Chicago. Chicago, IL USA; Department of Neurological Surgery, Feinberg School of Medicine, Northwestern University, Chicago, IL USA; Northwestern Medicine Malnati Brain Tumor Institute of the Lurie Comprehensive Cancer Center, Feinberg School of Medicine, Northwestern University; Department of Pathology, Feinberg School of Medicine, Northwestern University, Chicago, IL, USA; Department of Pediatrics, Icahn School of Medicine at Mount Sinai, New York, NY 10029

## Abstract

**BACKGROUND:** Diffuse midline gliomas remain incurable, with consistently poor outcomes in children despite radiotherapy. Immunotherapeutic approaches hold promise, with the integration of the host’s immune system fundamental to their design. Here, we describe a new, genetically engineered immunocompetent model that incorporates interleukin 13 receptor alpha 2 (IL13Rα2), a tumor-associated antigen, which is suitable for further evaluation of the antitumor activity of IL13Rα2-targeted immunotherapeutics in preclinical studies.

**METHODS:** The RCAS-Tv-a delivery system was used to induce gliomagenesis through overexpression of PDGFB and p53 deletion with and without human IL13Rα2 in Nestin-Tva; p53^fl/fl^ mice. Hindbrain or cerebral cortex nestin progenitors of neonatal pups were infected with Cre recombinase and PDGFB+IL13Rα2 or Cre recombinase and PDGFB to model diffuse midline glioma and supratentorial high-grade glioma, respectively. Immunoblotting and flow cytometry was used to confirm target expression. Kaplan-Meier survival curves were established to compare tumor latency in both models. Tumor tissue was analyzed through immunohistochemistry and H&E staining. Cell lines generated from tumor-bearing mice were used for *in vitro* studies and orthotopic injections.

**RESULTS:** The protein expression of PDGFB and IL13Rα2 was confirmed by flow cytometry and western blot. In both groups, *de novo* tumors developed without significant difference in median survival between PDGFB and p53 loss (n=25, 40 days) and PDGFB, IL13Rα2, and p53 loss (n=33, 38 days, p=0.62). Tumors demonstrated characteristics of high-grade glioma such as infiltration, palisading necrosis, microvascular proliferation, high Ki-67 index, heterogeneous IL13Rα2 expression, and CD11b+ macrophages, along with a low proportion of CD3+ T cells. Orthotopic tumors developed from cell lines retained histopathological characteristics of *de novo* tumors. Mice orthotopically implanted with cells in the hindbrain or right cortex showed a median survival of 42 days and 41 (p=0.56) days, respectively.

**CONCLUSION:** Generation of *de novo* tumors using the RCAS-Tv-a delivery system was successful, with tumors possessing histopathologic features common to pediatric diffuse gliomas. The development of these models opens the opportunity for preclinical assessment of IL13Rα2-directed immunotherapies with the potential for clinical translation.

## INTRODUCTION

Among pediatric cancers, diffuse midline gliomas are notorious for their universally poor outcomes^1^. The prototypical diffuse midline glioma, diffuse intrinsic pontine glioma (DIPG), is an aggressive tumor that has retained a median survival of less than one year from diagnosis and remains without long-term curative therapy^2^. Radiation provides limited symptom control, with an average extension of overall survival by three months. The lack of sustained response to conventional therapies reinforces the pressing need for new therapies.^3,4^

With recent achievements in cancer immunotherapy in adults and children, the push to develop immunomodulatory applications in the central nervous system (CNS) tumors intensifies. Advances in molecular characterization of midline gliomas, made through increased tissue sampling at diagnosis and post-mortem, have enabled subgrouping of tumors and identification of molecular drivers^1,5-7^. Surface markers, such as GD2 and B7-H3, have previously been characterized based on expression levels and degree of heterogeneity, both within tumors and amongst patients^8,9^. The preclinical studies validation of GD2 and B7-H3 as targets for chimeric antigen receptor-modified T cells (CAR T-cells) has led to recently initiated clinical trials (NCT04196413, NCT04099797, NCT04185038)^10,11^. Another attractive immunotherapeutic target in pediatric HGG is interleukin-13 receptor alpha 2 (IL13Rα2). High expression of IL13Rα2 has been described in DIPG and other pHGG tissues^4,12,13^. Within adult-type diffuse gliomas, IL13Rα2 targeting therapies have been the focus of multiple preclinical and clinical studies. There is currently one pilot study utilizing the IL13Rα2 epitope for a glioma vaccine^14^, featuring IL13Rα2 targeting CAR T-cells in IL13Rα2-positive brain tumors (NCT02208362). Within pediatric high-grade gliomas (pHGG), there is currently a phase one trial utilizing IL13Rα2 targeting autologous CAR T-cells in patients with recurrent or refractory brain tumors (NCT04510051). Although these advances provide a growing number of targets for immunotherapy and justify further investigation into IL13Rα2 as a target, the lack of an immunocompetent preclinical model of DIPG that expresses human IL13Rα2 hinders the development of IL13Rα2-directed immunotherapies.

One such system that permits tumorigenesis in the presence of a competent immune system is a genetically engineered mouse model (GEMM)^13,15^. GEMMs allow the introduction of specific genetic abnormalities in a spatiotemporal manner and the ability to study various stages of tumor development, including tumor initiation^16^. A versatile brainstem glioma GEMM was developed by Becher and colleagues using the RCAS/Tv-a system to initiate tumorigenesis in newborn mice^17^. This model allows for somatic cell gene transfer to cells engineered to express the Tv-a receptor^18^. By engineering Tv-a receptors to nestin-positive brainstem progenitor cells, a candidate cell of origin for DIPG^19,20^, it is possible to induce tumorigenesis to model DIPG. Furthermore, this highly adaptable model can alter the expression of genes of particular interest, namely immunotherapeutic targets, within Tv-a-expressing cells.

Using this RCAS/Tv-a system and integrating IL13Rα2 into the construct encoding for PDGFB, we have developed a novel immunocompetent GEMM that recapitulates diffuse midline gliomas while expressing human IL13Rα2. In doing so, we created a GEMM that permits the assessment of IL13Rα2-directed immunotherapeutic approaches for diffuse midline gliomas and pHGG. We have previously developed an IL13Rα2 specific antibody and have incorporated it into both CAR T-cells and bi-specific T-cell engagers for preclinical evaluation in adult glioblastoma.^21,22^ Given our experience, we are well-equipped to carry out studies in pediatric tumors. Engineered midline diffuse and high-grade glioma that express IL13Rα2 will fill an important gap in existing models and will allow preclinical mechanistic and therapeutic studies in corresponding pediatric brain tumors.

## MATERIALS and METHODS

### Analysis of TCGA patient data sets

Levels of IL13Rα2 gene expression in pediatric brain tumors were assessed using a publicly available microarray expression profile Gump database (tumor samples: 106, non-tumor: 6; GEO ID: GSE68015). RNA-seq data from Gump were visualized using GlioVis (http://gliovis.bioinfo.cnio.es/), with the relative expression of IL13Rα2 determined from log2 transformation of normalized count reads.^23,24^ Statistical significance between pediatric tumors was determined using Tukey’s Honest Significant Difference (HSD) within GlioVis, with one asterisk (*) where p<0.05, two (**) where p<0.01, and three (***) where p<0.001.

### Histology and Immunohistochemistry of human samples

Human tumor samples were initially obtained under Institutional Review Board approved protocol (#IRB 2019-3164) per Ann and Robert H. Lurie Children’s Hospital of Chicago. All samples were deidentified. Tissue sections were stained with H&E and for IL13Ra2 (R&D antibody, cat AF146) IHC. Scoring was performed independently by a neuropathologist (C.H.). Scores were established on percent positively labeled tumor cells; 0 is no positivity, 1 is ≤10% positive tumor cells, 2 is 10–50% positive tumor cells, 3 is ≥50% positive tumor cells.

### Cell lines and cell culture reagents

DF-1 (chicken fibroblast) cell line was obtained from ATCC (cat #CRL-12203). All cell lines were cultured in Dulbecco’s Modified Eagle’s Medium (DMEM) (ATCC, Manassas, Virginia; Cat #30-2002) supplemented with 10% fetal bovine serum (FBS) (R&D Systems, Minneapolis, MN), 1% penicillin/streptomycin (PS) (Corning, Corning, NY; Cat #30-002-CI) and 2mM L-glutamine (Corning, Corning, NY; Cat # 25-005-CI). Dulbecco’s Phosphate-buffered saline (PBS) was obtained from Thermo Fisher. GeneJuice was purchased from (Millipore Sigma).

### Cloning IL13Rα2 in vector encoding for PDGFB cDNA

The cDNA encoding T2A-IL13Ra2 was synthesized by GenScript, subcloned into the RCAS-PDGFB-HA vector^25^ following the PDGFB cDNA using CloneEZ method, and verified by Sanger sequencing (GenScript, Piscataway, NJ). The plasmid was amplified in DH5a cells (NewEngland Biolabs, Ipswich, MA; Cat #C2987H), and purified using plasmid purification midi-kit (Qiagen, Hilden, Germany; Cat # 12145).

### Generation of DF-1 cells producing virus encoding for PDGFB, RCAS-Cre, and PDGFB+IL13Rα2

Our approach expands upon a previously described use of the RCAS-Tva system for GEMM creation.^14^ Briefly, we cultured DF-1 cells in optimal growth conditions in DMEM supplemented with 10% FBS, 1% PS, and 2 mM L-glutamine. Cells were transfected with RCAS plasmids (RCAS-Cre, PDGFB, and PDGFB+IL13Rα) using GeneJuice (Millipore Sigma, Burlington, MA; Cat # 70967), per the manufacturer’s protocol. In order to propagate the necessary cell number and adequate retrovirus for intracranial injection, we serially passaged the three transfected cell groups upon reaching 80-90% density for up to five passages.

### Western blot

Cells were cultured in complete DMEM as previously described. Cells were collected, pelleted, and then resuspended into one mL of cold Dulbecco’s Phosphate-buffered saline (PBS) (4°C). Cells were counted via the trypan-blue exclusion method, pelleted by centrifugation (2000 x g, 5 min), and lysed in ice-cold Mammalian Protein Extraction Reagent (mPER) (Thermo Scientific, Waltham, MC) at a concentration of 20,000 cells/μL mPER. Protein concentration was measured using a PierceTM bicinchoninic acid (BCA) protein assay kit (Thermo Scientific) per manufacturer recommendations. Protein samples were then prepared in Laemmli Sample Buffer (Bio-Rad, Hercules, CA; Cat #1610747) with 2.5% 2-Mercaptoethanol (Sigma Aldrich, St. Louis, MO; Cat #M6250) and denatured at 95 °C for 5 min. Protein samples and 250 kDa Precision Plus Protein standard (Bio-Rad Laboratories, Hercules, CA) were loaded into each well of a 4-20% Midi-PROTEAN® TGX Stain-Free™ gel (Bio-Rad). Proteins were separated by size using SDS-polyacrylamide gel electrophoresis (SDS-PAGE) and transferred onto a methanol-activated 0.2 μm PVDF membrane (Millipore Sigma, Burlington, MA; Cat #IPVH08100) using the Trans-Blot® Turbo Transfer System™ (Bio-Rad). Membranes were blocked with 5% non-fat dry milk (NFDM) in tris-buffered saline plus 0.1% Tween-20 (TBST) (Boston Bioproducts, Ashland, MA; Cat #IBB180) for 2 hours at RT. Membranes were incubated with anti-IL13Rα2 Monoclonal Antibody (1:1000) (R&D Systems, AF146), anti-HA tag antibody to detect PDGFB (1:1000) (Cell Signaling Technology, 3724), and anti-glyceraldehyde 3-phosphate dehydrogenase (GAPDH) Monoclonal Antibody (1:1000) (Cell Signaling Technology, 2118) in 5% NFDM-TBST overnight at 4°C. The next day, membranes were washed with TBST and incubated with anti-rabbit HRP-linked Secondary Antibody (1:1000) (Cell Signaling Technology, 7074) for GAPDH and anti-HA tag and chicken anti-goat Secondary Antibody (Santa Cruz Biotechnology, sc-2953) for IL13Rα2. Blots were washed with TBST, incubated with Clarity™ Western ECL Substrate (Bio-Rad), and detected via chemiluminescent image acquired by ChemiDoc™ MP Imaging System (Bio-Rad).

### Flow Cytometry

Cells were maintained in culture with complete DMEM. Cells were collected, pelleted, and incubated with TruStain Fc Blocking Antibody (1:200 in 1% FBS in DPBS) (4°C). Next, 100,000 cells were added to each well of a TC treated V-bottom plate. Cells were pelleted and stained on ice with PE anti-human IL13Rα2 (Biolegend, 354403). Live cells were labeled with a fixable viability stain (Invitrogen, 65-0865-14). Cells were then fixed, permeabilized, and stained with Alexa-Fluor 488 Anti-HA antibody (Biolegend, 901509) to detect PDGFB. Data acquisition was done using the FACSymphony A5-Laser Analyzer, BD.

### Histology and Immunohistochemistry of mouse samples

At selected time points or based on clinical deterioration, animals were perfused with PBS to collect brain tissue for sectioning. Tissue was fixed in 4% paraformaldehyde (PFA), then paraffin-embedded, cut into 4 microns sections, and stained with hematoxylin and eosin (H&E). Further immunohistochemistry staining was obtained using following antibodies: IL13Rα2 (R&D AF146; 1:200), Olig2 (Abcam ab109186; 1:2000), Ki67 (Abcam ab16667; 1:500) CD3 (Abcam ab16669; 1:1000) CD11b (Abcam ab133357; 1:3000).

### Animal Studies

Nestin-tva (Ntv-a) p53^fl/fl^ mice were bred in-house per animal protocols approved by the Northwestern University Institutional Animal Care and Use Committee (IACUC). Suspensions of cells producing virus were injected into the hindbrain of Ntv-a p53^fl/fl^ mice pups, between three to five post-natal days of age, using cryoanesthesia during the procedure. We evaluated two experimental groups: 1) RCAS-PDGFB, and 2) RCAS-PDGFB+IL13Rα2, with the former as the established model and the latter our new model with a combined cell count of 2×10^5^ cells/mL per pup (a 1:1 ratio of RCAS-Cre with PDGFB or PDGFB+IL13Rα2) suspended in PBS for a total volume of 1.2 µL. The injection site is approximately 2 mm posterior to the bregma along the midline using a Hamilton syringe and custom needle. After postprocedural recovery and rewarming, pups were returned to their nursing mother. Following injection, mice were monitored closely for symptoms consistent with tumor involvement, including ataxia, seizures, enlarged head, and weight loss.

### Generation of cell lines for orthotopic model

Cell lines were generated by processing tumor-bearing tissue obtained from symptomatic mice. Mice were anesthetized with a solution consisting of 200 mg/kg ketamine and 20 mg/kg xylazine and perfused with PBS before neoplastic tissue was processed with the NeuroCult™ enzymatic dissociation Kit (StemCell Technologies, 05715) per manufacturer’s instructions. Cells were maintained as neurospheres in NeuroCult™ proliferation media (Stem Cell Technologies, Vancouver, Canada; Cat #05072) with the addition of mouse cell proliferation supplement (Stem Cell Technologies, Cat #05701), PS, heparin at 2μg/mL and 10ng/mL epidermal growth factor, and 20ng/mL basic fibroblast growth factor. For orthotopic experiments, mouse pups were injected with 1 × 10^5^ cells (1.2 µL) per animal in the hindbrain and the right cortex, as described in animal studies.

### Statistics

Statistical analyses of data were executed using GraphPad 8 (Prism, La Jolla, CA) and Microsoft Excel. Significance was defined as *p* less than 0.05 in all statistical tests. P values were established as follows: \**p* < .05, \*\**p* < .01, *** *p* < .001, and **** *p* < .0001. As indicated, data in two groups were analyzed for statistical significance using the unpaired Mann-Whitney test or an unpaired two-tailed Student’s t-test. One- or two-way ANOVA was used for multiple groups, followed by a Tukey’s or Dunnett’s multiple comparisons test. Animal survival analysis was performed by generating Kaplan-Meier plots using the log-rank method to determine the p-value.

## RESULTS

### Analysis of IL13Rα2 expression in patient tissues

IL13Rα2 expression in adult-type diffuse high-grade gliomas is well established, and it has been used as an immunotherapeutic target in multiple preclinical and clinical studies.^21,22,26,27^ To determine the relative expression in pediatric CNS tumors, we used GlioVis to visualize IL13Rα2 gene expression within the publicly available Gump database (GEO ID: GSE68015). Increased IL13Rα2 mRNA expression was found in pediatric-type diffuse high-grade gliomas (pHGG) compared to non-neoplastic brain tissue (Fig. 1A). Furthermore, elevated IL13Rα2 gene expression was seen in pHGG compared to low-grade CNS tumors (Fig. 1A). To further validate these findings, we assessed tissue expression of IL13Rα2 in twenty post-mortem pediatric DIPG samples using immunohistochemistry (IHC). Nine of the twenty (45%) samples were positive for IL13Rα2, receiving histopathological scores of 1 (n=5), 2 (n=3), and 3 (n=1) (Fig. 1B&C).

**Figure 1.**
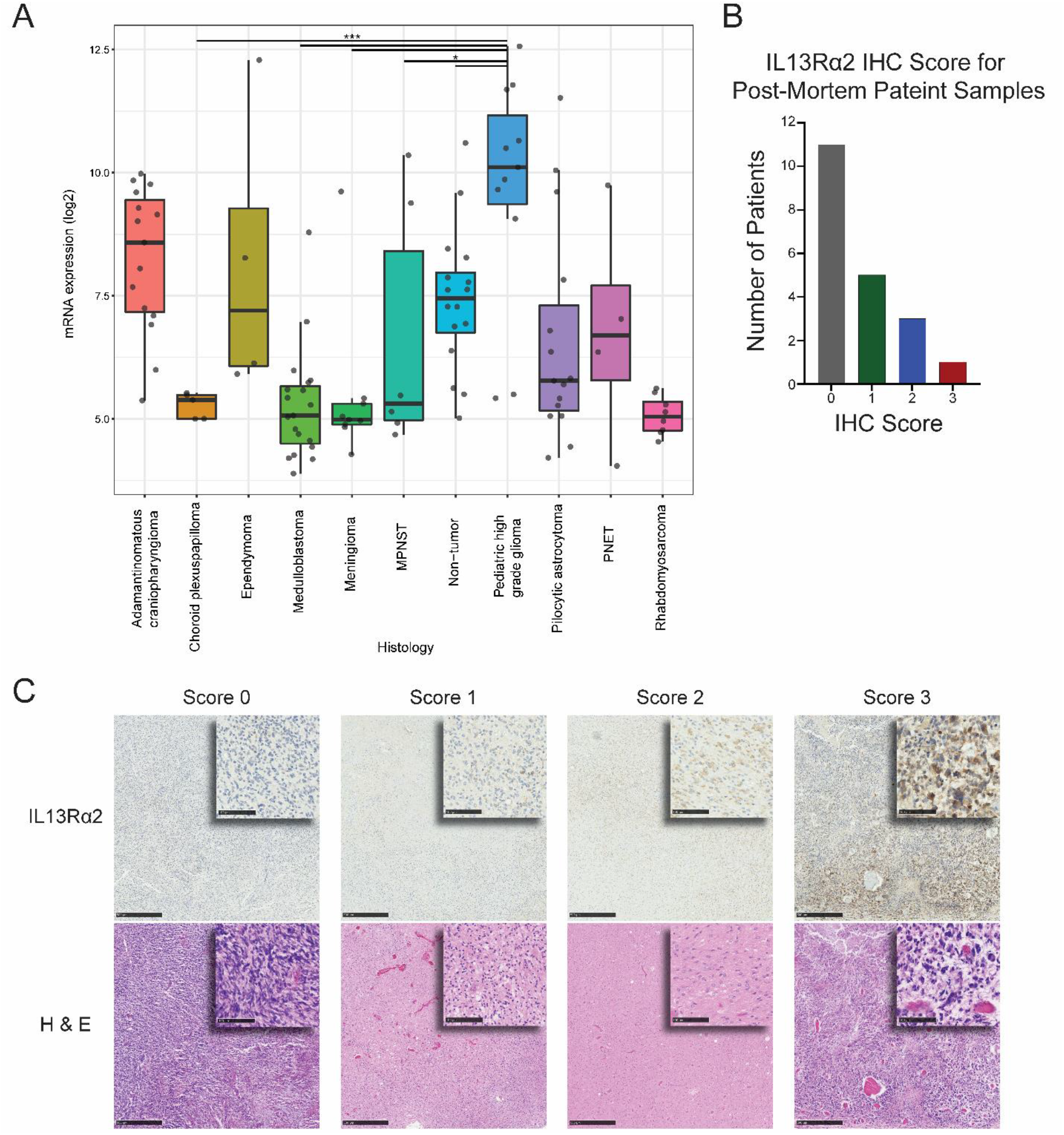
Heterogeneity of IL13Rα2 expression in human DIPG. (A) IL13Rα2 gene expression in pediatric brain tumor from Gump database was analyzed using GlioVis (http://gliovis.bioinfo.cnio.es/). Tukey’s Honest Significant Difference (HSD) was performed to compare IL13Rα2 gene expression between pediatric tumors. Significance was denoted in the graph by one asterisk (*) where p<0.05, two (**) where p<0.01, and three (***) where p<0.001. (B) IL13Rα2 IHC score of patient samples obtained post mortem. (C) Analysis of IL13Rα2 expression by IHC in post-mortem human DIPG patient samples (top panel), along with corresponding H&E stains (bottom panel). The degree of IL13Rα2 expression was scored following IHC test scoring method. High-power insets are derived from low-power fields.

### Generation and characterization of the RCAS-Tva model of high-grade pediatric diffuse glioma expressing IL13Rα2

To generate *de novo* tumors, we first cloned human IL13Rα2 in a PDGFB construct using a T2A cleaving peptide. Next, we transfected DF-1 cells with either RCAS-Cre, RCAS-PDGFB, or RCAS-PDGFB+IL13Rα2 constructs. Subsequent flow cytometry (FSC) analysis of DF-1 cells showed robust expression of PDGFB (93.8%) and PDGFB+IL13Rα2 (73.5%) (Fig. 2B). Secondary confirmation of successful RCAS-PDGFB, and RCAS-PDGFB+IL13Rα2 transfection was achieved via western blotting for PDGFB and IL13Rα2 (Fig. 2C). Following confirmation of transduction, we evaluated growth dynamics of de novo tumors *in vivo* by infecting nestin-expressing progenitors in either the hindbrain or right cortex of 4-5 day-old Nestin-*tva* p53^fl/fl^ mice to model pediatric diffuse midline glioma (pDMG) and pHGG, respectively. No significant changes were noted in median survival of mice between RCAS-PDGFB (n=25, 40 days) and RCAS-PDGFB+IL13Rα2 (n=33, 38 days) of pDMG models (Fig. 2D). The median survival of mice injected in the right cortex with the RCAS-PDGFB+IL13Rα2 system, a model pHGG, was 45.5 days (2E).

**Figure 2.**
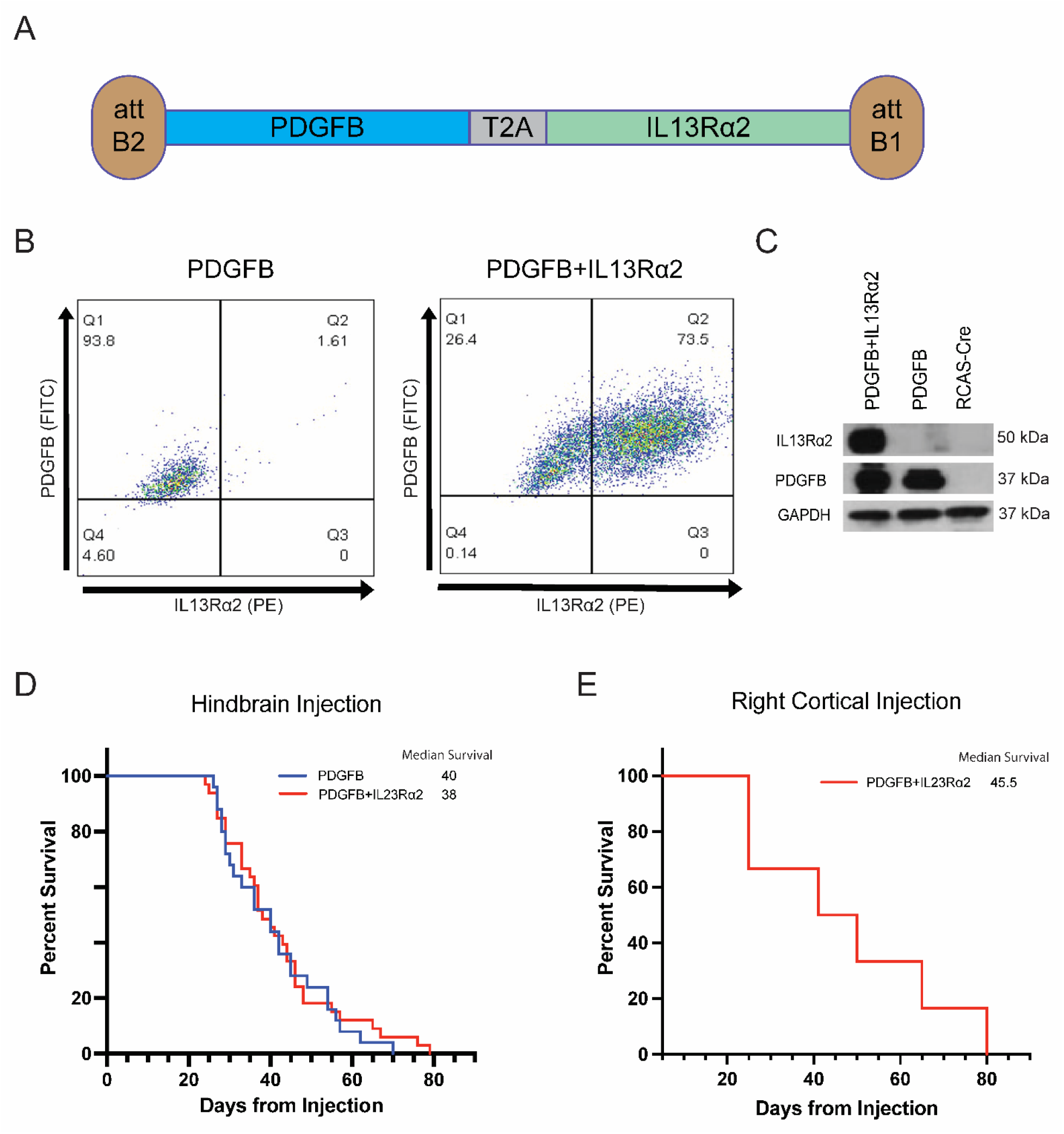
Analysis of GEMM models of DIPG and midline glioma expressing IL13Rα2. (A) Schematic of Tva vector generated to encode PDGFB andIL13Rα2 transgenes and used to transfect DF-1 cells. Successful expression of PDGFB and IL13Rα2 in DF-1 was confirmed by (B) flow cytometry and (C) western blot. (D) Survival curves of Nestin-Tva;p53^fl/fl^ mice injected with DF-1 producing RCAS-PDGFB and RCAS-PDGFB+IL13Rα2 in the hindbrain, and (E) RCAS:PDGFB+IL13Rα2 in the cortical area of the brain.

### Histopathological analysis of de novo tumors

To further evaluate the recapitulation of our GEMM model to pediatric tumors in humans, we performed IHC analysis for various markers. Both pDMG models (Fig 3A) and pHGG models (Fig. 3B) revealed similar characteristics to post-mortem patient samples,^1^ with noticeable cellular atypia, microvascular proliferation, and pseudopalisading necrosis in *de novo* tumors (Fig. 3A and B). Additional characteristics of DIPG were present in both GEMM models, with a cellular invasion of adjacent parenchyma, Ki67 positivity, Olig-2 expression, the presence of CD11b+ cells, low number of CD3+ cell, and heterogeneous IL13Rα2 expression (Fig. 3A and B).

**Figure 3.**
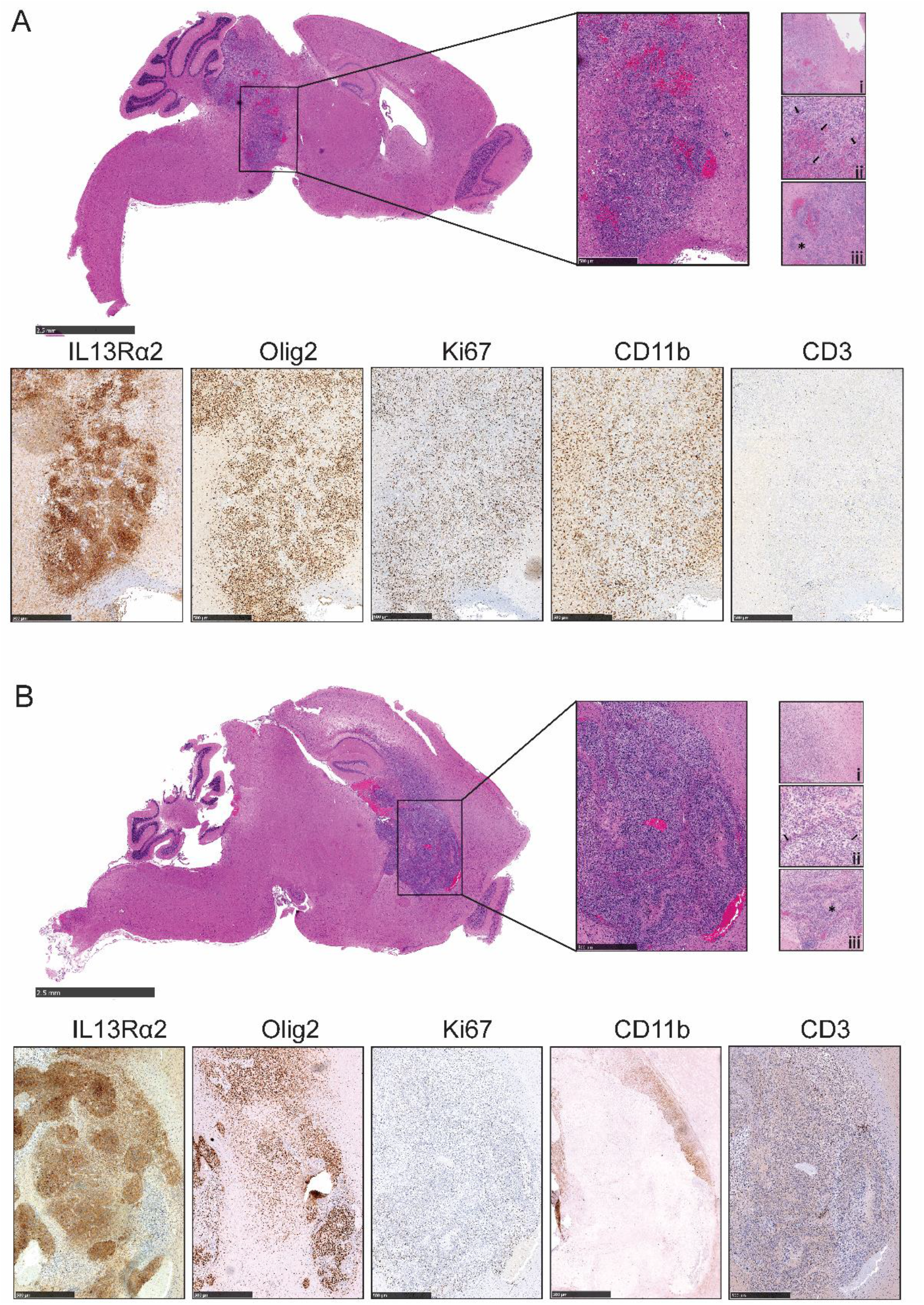
Histopathological analysis of *de novo* tumors. (A) *De novo* tumor generation for diffuse midline glioma through hindbrain injection, 49 days post-injection. Low magnification of whole brain tissue section and tumor stained with H&E. Higher magnification of inset images reveals infiltrative tumor cells (i), microvascular proliferation (ii, arrows), and pseudo-palisading necrosis (iii, asterisk). (B) Similar characterization of *de novo* tumor generation of supratentorial high-grade glioma through right cortex injection, 50 days post-injection.

### Generation of cell lines and orthotopic transplantation

In order to develop targeted immunotherapies using these models, we generated cell lines from our *de novo* tumors. Brain tissue from tumor-bearing animals was processed in a single-cell suspension as described in the Material and Methods, placed into culture, and maintained as neurospheres. Expression of PDGFB and IL13Rα2 was confirmed by FSC analysis and western blot analysis (Fig. 4A and B). Next, cell lines were orthotopically implanted into the hindbrain ventricle or right cortex of 4-5 days-old N-tva p53^fl/fl^ mice. Survival analysis displayed a median survival of 42 and 41 days for mice injected in the hindbrain and right cortex, respectively (Fig. 4C). IHC analysis of these tumors displayed histological features similar to those of *de novo* tumors, with heterogeneous IL13Rα2 expression, Olig-2 expression, infiltration of CD11b+ cells, and a small number of CD3+ cells along with Ki67-positivity (Fig. 4D).

**Figure 4.**
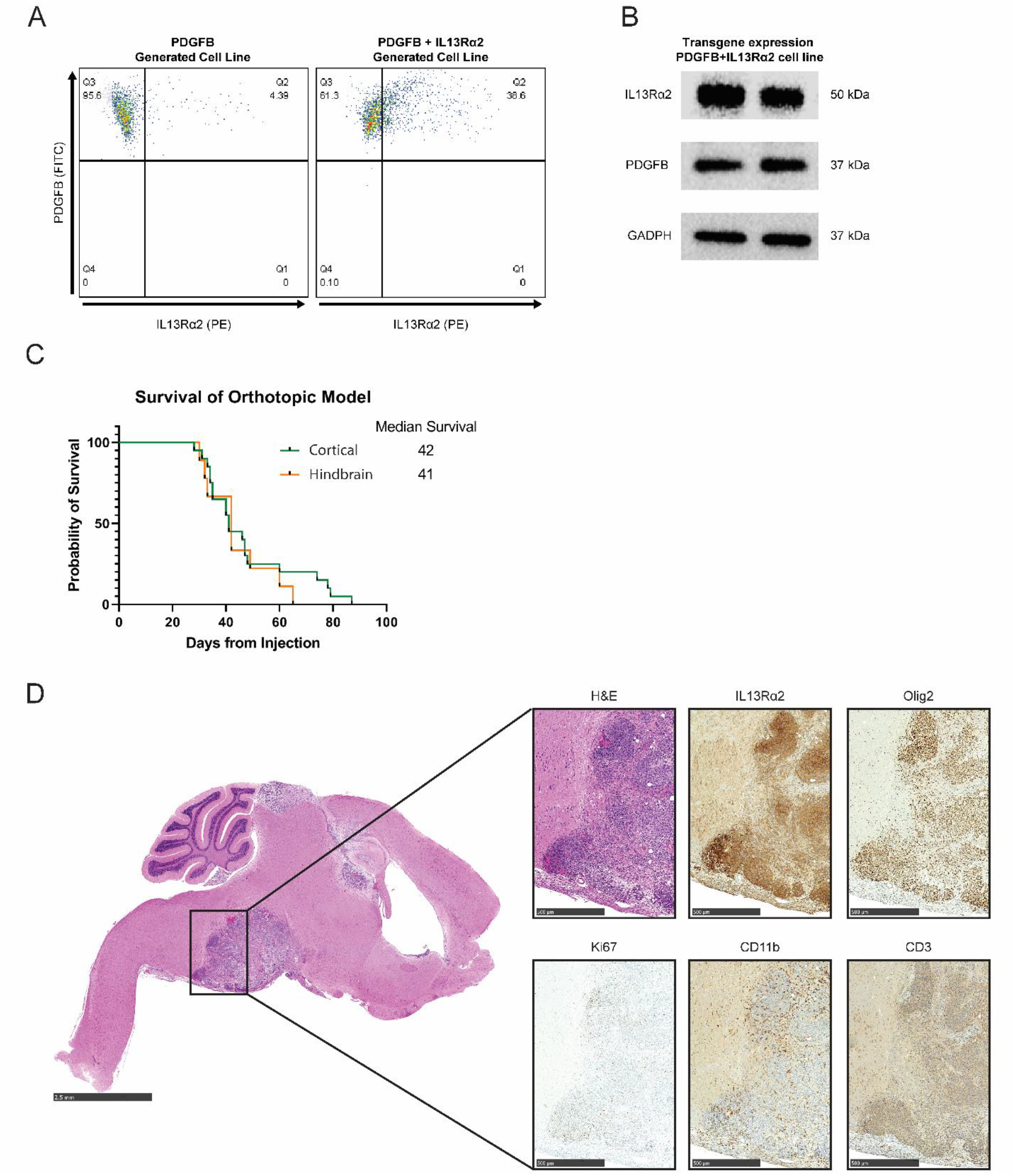
Generation and analysis of cell lines generated from *de novo* tumors. (A) Flow cytometry analysis of IL13Rα2 and PDGFB expression in cell lines generated from *de novo* tumors. (B) Secondary confirmation of PDGFB and IL13Rα2 expression in cell lines by western blot. (C) Survival analysis of orthotopically injected cell lines in the hindbrain and cortex of Nestin-Tva; p53^fl/fl^ mice. (D) Histopathological analysis of orthotopically injected cell lines, with staining for IL13Rα2, Olig2, Ki67, CD11b, and CD3.

## DISCUSSION

Among many other factors, the efficacy of targeted immunotherapy is dependent on the validity of the targeted antigen. IL13Rα2 is an attractive TAA present in several CNS tumors, and it has been validated as an immunotherapeutic target in adult GBM^28,29^. To better understand the potential of IL13Rα2 as an immunotherapeutic target in the context of pHGG, we first characterized its expression in datasets of patient tissue samples. We found that mRNA for IL13Rα2 is expressed at higher levels in pHGG, and about that 45% of DIPG samples expressed IL13Rα2, in agreement with other reports.^8,13^. Thus, IL13Rα2 has the potential to be used as an immunotherapeutic target for DIPG and pHGG.

A major obstacle to the development of targeted immunotherapy for pDMG and pHGG is the lack of an immunocompetent model expressing the corresponding antigen. We used the RCAS-Tv-a system to induce *de novo* tumorigenesis in immunocompetent mice to overcome this limitation. In doing so, we were able to recreate diffuse midline glioma and high-grade glioma that express IL13Rα2. We demonstrate that the addition of IL13Rα2 to PDGFB-driven gliomas did not significantly impact the median survival of the mice in all experimental settings compared to the previously described PDGFB-driven tumor model of pediatric diffuse midline glioma^30^. Histopathologic characteristics of *de novo* tumors were similar to those seen in patient samples, such as cellular atypia, microvascular proliferation, pseudopalisading necrosis, and invasion into surrounding brain tissue.

Generated cell lines expressed transgene products of the initial constructs and reliably generated tumors when implanted orthotopically in the murine brain. Both *de novo* and orthotopically generated tumors have a heterogeneous expression of IL13Rα2, consistent with that found in pediatric diffuse midline gliomas and GBM. Consistent with T-cell infiltration described in pediatric gliomas ^31,32,25^, IHC analysis of tumor microenvironment showed limited and scattered CD3 positive cells and a moderate presence of CD11b+ cells in both *de novo* and orthotopic diffuse midline tumors. Further analysis of the tumor microenvironment will be pursued to understand these findings better and to assess changes taking place upon the introduction of immunotherapy.

One limitation of this work is that we did not include the H3.3K27M in the model, which is expressed in most diffuse midline gliomas^5,6^. However, this work is a proof of concept that relevant human antigens can be readily incorporated into GEMMs and can serve as useful preclinical tools. Future work will also incorporate the H3.3K27M onco-histone as previously reported^30^.

We envision that the developed models will permit for preclinical assessment of IL13Rα2-targeting immunotherapies, including CAR T-cell, BiTE, and bispecific killer cell engagers (BiKE) therapies within an immunocompetent host. Upon establishing the potential antitumor effect of each of these therapies, combinatorial approaches either with concomitant radiation, medications such as chemotherapy or targeted therapies, or in conjunction with other immunotherapeutic approaches such as oncolytic viruses, checkpoint inhibitors, or other antibodies will then be considered. A robust evaluation of such approaches will allow for potential clinical translation to improve the outcome for children impacted by these devastating tumors.

## ACKNOWLEDGEMENTS

This study was supported in part by NINDS R01NS106379 (IVB) and R01NS122395 (IVB) grants and the Washington Square Health Foundation (MS). Histology services were provided by the Northwestern University Mouse Histology and Phenotyping Laboratory, which is supported by National Cancer Institute Grant P30-CA060553 awarded to the Robert H. Lurie Comprehensive Cancer Center. In addition, this work was supported by the Northwestern University – Flow Cytometry Core Facility supported by Cancer Center Support Grant (NCI CA060553). The Northwestern Nervous System Tumor Bank is supported by the P50CA221747 SPORE for Translational Approaches to Brain Cancer. The authors are thankful to Liliana Ilut for her technical assistance on this project.

